# Structural basis of asymmetric transcription through a composite nucleosome formed by a hexasome and an octasome

**DOI:** 10.64898/2025.12.26.696134

**Authors:** Zhihui Chen, Cheng-Han Ho, Hiroki Tanaka, Tomoya Kujirai, Mitsuo Ogasawara, Haruhiko Ehara, Shun-ichi Sekine, Yoshimasa Takizawa, Hitoshi Kurumizaka

**Author notes:** Present address: Department of Structural Virology, National Center for Global Health and Medicine, 1-21-1 Toyama, Shinjuku-ku, Tokyo, 162-8655, Japan (Hiroki Tanaka). These authors contributed equally to this work (Zhihui Chen, Cheng-Han Ho).

## Abstract

The overlapping dinucleosome (OLDN), a composite chromatin particle consisting of a hexasome and a canonical octasome, forms immediately downstream of transcription start sites, likely through chromatin remodeling activity, and has been proposed to act as a transient regulatory intermediate during transcription. However, how RNA polymerase II (RNAPII) engages with and transcribes through this unusual structure remains unclear. Here, we reconstituted OLDNs *in vitro* and performed transcription assays with RNAPII. We found that transcription efficiency was markedly higher when RNAPII initiated from the hexasome side than from the octasome side. Cryo-electron microscopy further revealed that transcription from the hexasome side induced pronounced conformational rearrangements, in which RNAPII progression dramatically opened the hexasome-octasome interface. These results uncover the mechanism by which RNAPII senses the intrinsic transcriptional polarity of OLDNs and suggest that OLDNs function as dynamic, directionally sensitive regulators of transcription elongation.

## Main text

In eukaryotic cells, genomic DNA is packed into the cell nucleus by the formation of chromatin, which poses a substantial barrier to DNA-templated processes such as transcription^1–5^. The nucleosome, composed of ∼147 base pairs (bps) of DNA wrapped around the histone octamer containing two H2A-H2B dimers and one (H3-H4)_2_ tetramer, serves as the fundamental unit of chromatin^6^. In the nucleosome, the DNA is symmetrically wrapped around the histone octamer with a periodicity of approximately 10 base pairs, defining discrete positions known as superhelical locations (SHLs). The SHL(0) position marks the DNA center located at the dyad axis of the nucleosome, and positive and negative SHL values are assigned to the DNA superhelical turns on either side of the dyad ^6,7^. In chromatin, nucleosomes are connected by short linker DNAs, forming a “beads-on-a-string” conformation^8^. The nucleosome architecture must be dynamically remodeled to enable transcription by RNA polymerase II (RNAPII), which actively promotes nucleosome disassembly and reassembly during transcription elongation^9^.

A non-canonical chromatin structure termed the overlapping dinucleosome (OLDN) has been proposed as a potential product by transcription-associated chromatin remodeling factors, such as esBAF and ISWI, which slide nucleosomes along DNA to generate higher-order packing and transient structural intermediates ^10–12^. The crystal structure of the OLDN reveals that a hexasome lacking one H2A-H2B dimer is tightly associated with a neighboring canonical nucleosome (octasome)^13^. This unique dinucleosomal assembly wraps ∼250 bps of DNA without intervening linker DNA, resulting in a highly compacted chromatin configuration. Genome-wide analyses of ∼250-bp DNA fragments protected from extensive micrococcal nuclease digestion reveal that OLDNs are enriched immediately downstream of transcription start sites, corresponding to +1 nucleosome positions, suggesting that they are encountered frequently during transcription initiation and elongation ^13,14^. These pieces of evidence suggested that OLDNs are not merely passive obstacles to transcription, but may represent dynamic, chromatin-based regulators of gene expression.

The OLDN may also be formed when RNA polymerase II (RNAPII) promotes nucleosome disassembly and reassembly during transcription elongation, where a hexasome is formed as an intermediate^9,15^, and chromatin remodeler may slide the hexasome and collide it to adjacent nucleosome^16^. In addition, the OLDN may be observed in isolated mitotic chromosomes by a cryo-electron microscopy analysis^17^, suggesting that it persists throughout the cell cycle. However, the mechanism by which RNAPII engages and transcribes through OLDNs has remained unknown.

## Results

### Asymmetric RNAPII transcription in the overlapping dinucleosome

To test whether RNAPII can traverse the OLDN formed immediately downstream of transcription start sites, we first performed the RNAPII transcription assay with reconstituted OLDNs. We generated two derivatives depending on the orientation of the hexasome and octasome units relative to the transcription start site. To do so, an OLDN was assembled on a 250-bp tandem Widom 601 sequence, and a 73-bp linker DNA contained a 9-base mismatched region for the annealing site of a primer RNA and was ligated to a DNA end of the hexasome or octasome moiety (Fig.1a). These OLDN substrates allowed RNAPII transcription from either the hexasome or octasome side.

The reconstituted OLDNs were purified by a polyacrylamide gel electrophoresis. RNAPII was purified from the yeast *Komagataella phaffii* (formerly *Komagataella pastoris* or *Pichia pastoris*), and the elongation factor *K. phaffii* TFIIS was prepared as a recombinant protein^18,19^.

Our OLDN transcription assay revealed that RNAPII travers the OLDN from either the hexasome or octasome side (Fig.1b, c). Interestingly, RNAPII produced significantly more run-off transcripts when transcribing from the hexasome side than from the octasome side (Fig.1d). On naked DNA templates, the difference was much smaller, albeit still present. This suggests the asymmetric character of the OLDN in RNAPII transcription, which allows transcription more efficiently from the hexasome side than the octasome side. In addition, a specific RNAPII pausing at superhelical location (SHL) -0.5 position of the hexasome moiety (-5 bp position from the hexasome SHL(0) positon) was detected when the OLDN was transcribed from the hexasome side (Fig.1a, b). On the other hand, transcription pausing sites from the octasome side largely resemble those of the canonical nucleosome, in which RNAPII dominantly pauses at the SHL(-1) and SHL(0) positions of the octasome moiety (Fig.1a, c).

### The cryo-EM structures of RNAPII transcribing on the OLDN

To understand the mechanism by which the RNAPII traverses the OLDN from the hexasome side, we optimized the OLDN substrates and conducted the RNAPII transcription reaction in presence of either NTPs or ATP substituted with 3’deoxy-ATPs (see Methods). We determined two cryo-EM structures of the RNAPII-OLDN complex, RNAPII-OLDN^SHL(−6)^, and RNAPII-OLDN^SHL(−0.5)^, in which the leading edges of the RNAPII reached at the SHL(−6) and (−0.5) position of the hexasome moiety, respectively (Fig. 2a). These are structures of natural RNAPII pausing positions on the OLDN transcribed from the hexasome side. In the RNAPII-OLDN^SHL(−6)^ structure, the OLDN remains largely intact, in which the hexasome and octasome units are tightly associated, and closely resembles previously reported OLDN structures^13,20^.

**Figure 1.**
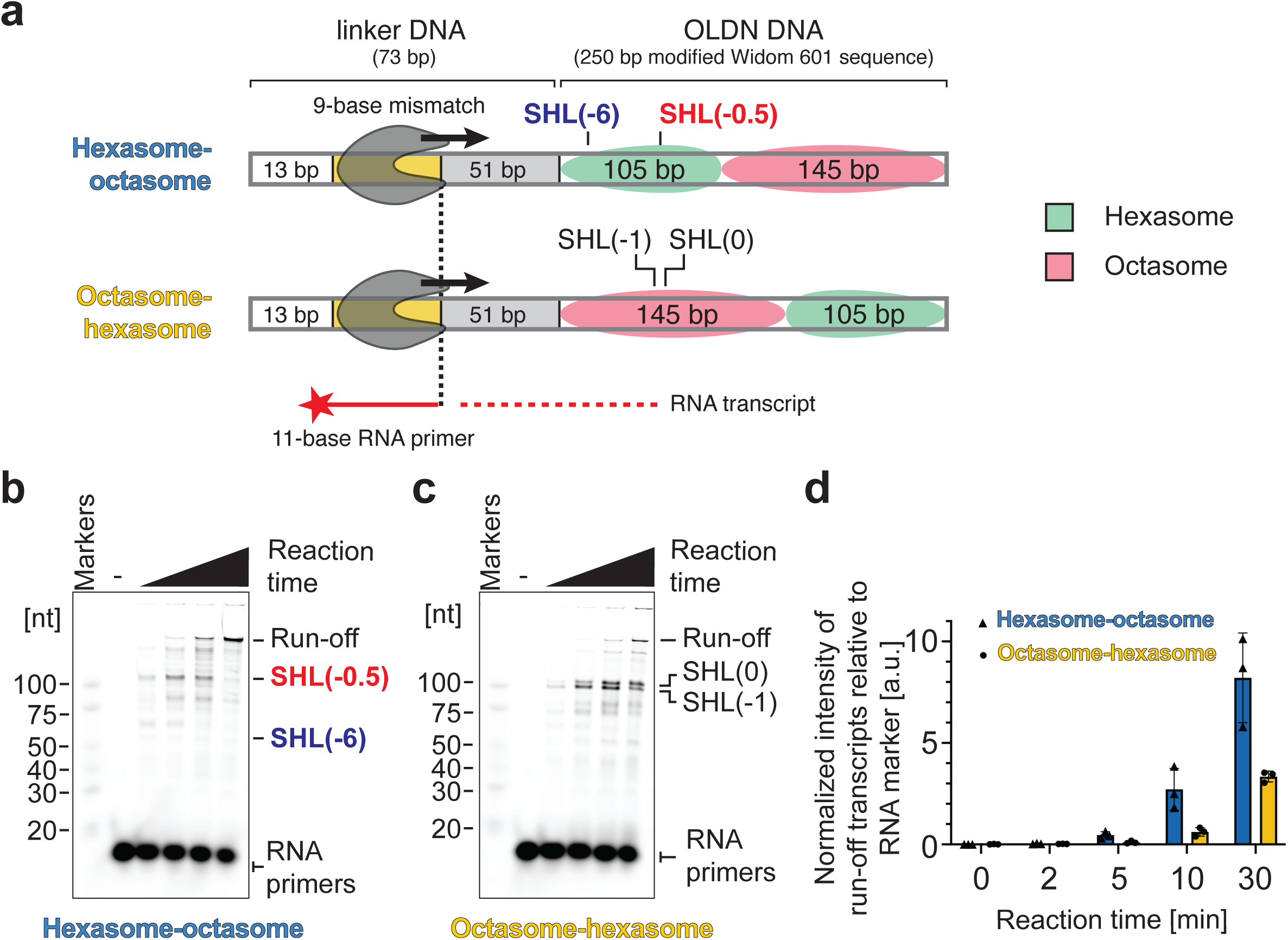
RNAPII traverses OLDN templates with different efficiencies depending on the entry side. **a,** Schematic representation of OLDN templates used for transcription assays. In the hexasome-octasome configuration (upper panel), RNAPII encounters the hexasome first, whereas in the octasome–hexasome configuration (lower panel), RNAPII encounters the octasome first. **b, c,** Representative transcription assay of the hexasome-octasome (**b**) and octasome-hexasome (**c**) configurations, cropped from the same gel. Assays were performed as time courses, with reaction mixtures collected at 0, 2, 5, 10 and 30 min and analyzed by denaturing PAGE. RNA transcripts were detected. Reproducibility of the transcription patterns was confirmed across three independent replicates. **d,** Quantification of run-off RNA transcripts from the hexasome-octasome and octasome-hexasome configurations across three independent experiments. Run-of RNA transcript levels were normalized to a reference ladder band on the same gel. Error bars indicate standard deviation (n = 3).

**Figure 2.**
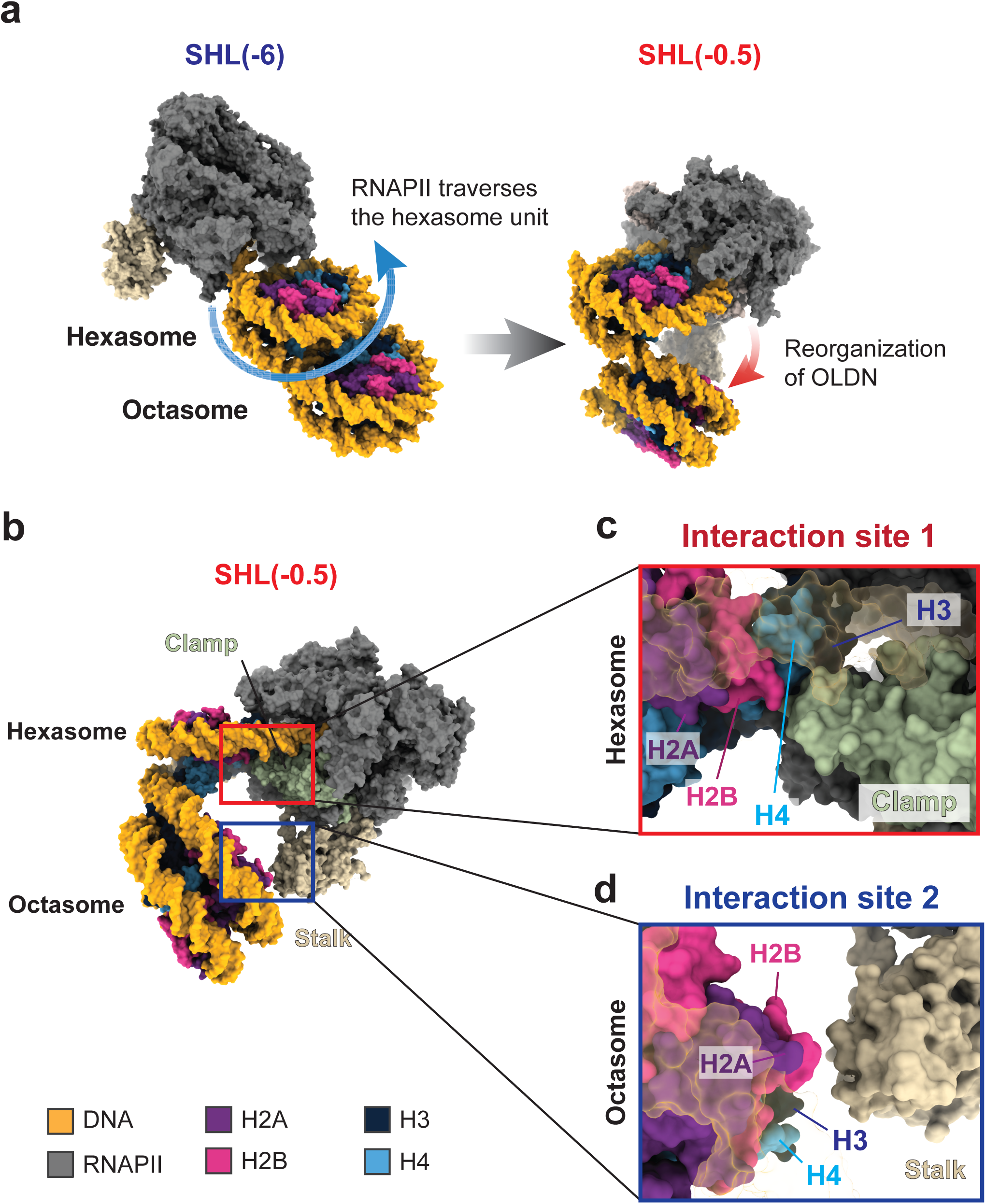
RNAPII induces extensive reorganization of OLDN subunits during transcription from the hexasome side. **a, Cryo-EM structures of the RNAPII-OLDN^SHL(−6)^ and RNAPII-OLDN^SHL(−0.5)^ complexes** formed during transcribed from the hexasome side. The two structures are aligned by the hexasome unit. **b,** Side view of the RNAPII-OLDN^SHL(−0.5)^ complex. RNAPII interacts with OLDN hexasome and octasome units at interaction site 1 and interaction site 2, respectively. **c,** The RNAPII clamp domain (Rpb1 subunit) directly engages H3, H4 and H2B within the hexasome moiety of the OLDN. DNA density in the foreground is rendered semi-transparent to visualize the histones. **d,** The RNAPII stalk domain interacts with the octasome moiety of the OLDN, potentially promoting opening or rearrangement of the OLDN structure.

Importantly, in the RNAPII-OLDN^SHL(−0.5)^ structure, we found that the overall conformation of OLDN is drastically changed; the hexasome and octasome moieties of OLDN are opened, with a nearly perpendicular angle between hexasome and octasome (Fig.2b). RNAPII Rpb1 clamp interacts with the distal surface of the hexasome unit, widening the hexasome-octasome interface (Fig. 2b, c). Concomitantly, the RNAPII subunit Rpb4-7 stalk directly contacts with the octasome moiety and pushes it away from the hexasome moiety (Fig.2b, d). The RNAPII-OLDN^SHL(−0.5)^ structure may represent a dynamic intermediate state, in which RNAPII actively remodels the OLDN configuration to facilitate RNA elongation.

### The template DNA looping may pause RNAPII at SHL(−0.5) of the hexasome unit

Interestingly, in the RNAPII-OLDN^SHL(−0.5)^ structure, the transcribed DNA region (upstream DNA) looped back to the remaining H2A-H2B of the hexasome moiety, forming the DNA looping conformation (Fig. 3a). This upstream DNA looping may be a major reason why the RNAPII paused at the SHL(−0.5) position. In the OLDN transcription assay, the RNAPII pausing at the hexasomal SHL(−0.5) position was detected and gradually decreased in the reaction time-dependent manner (Fig. 1b). Therefore, the upstream DNA looping with the hexasome may be transiently formed during the RNAPII traverse on the OLDN.

**Figure 3.**
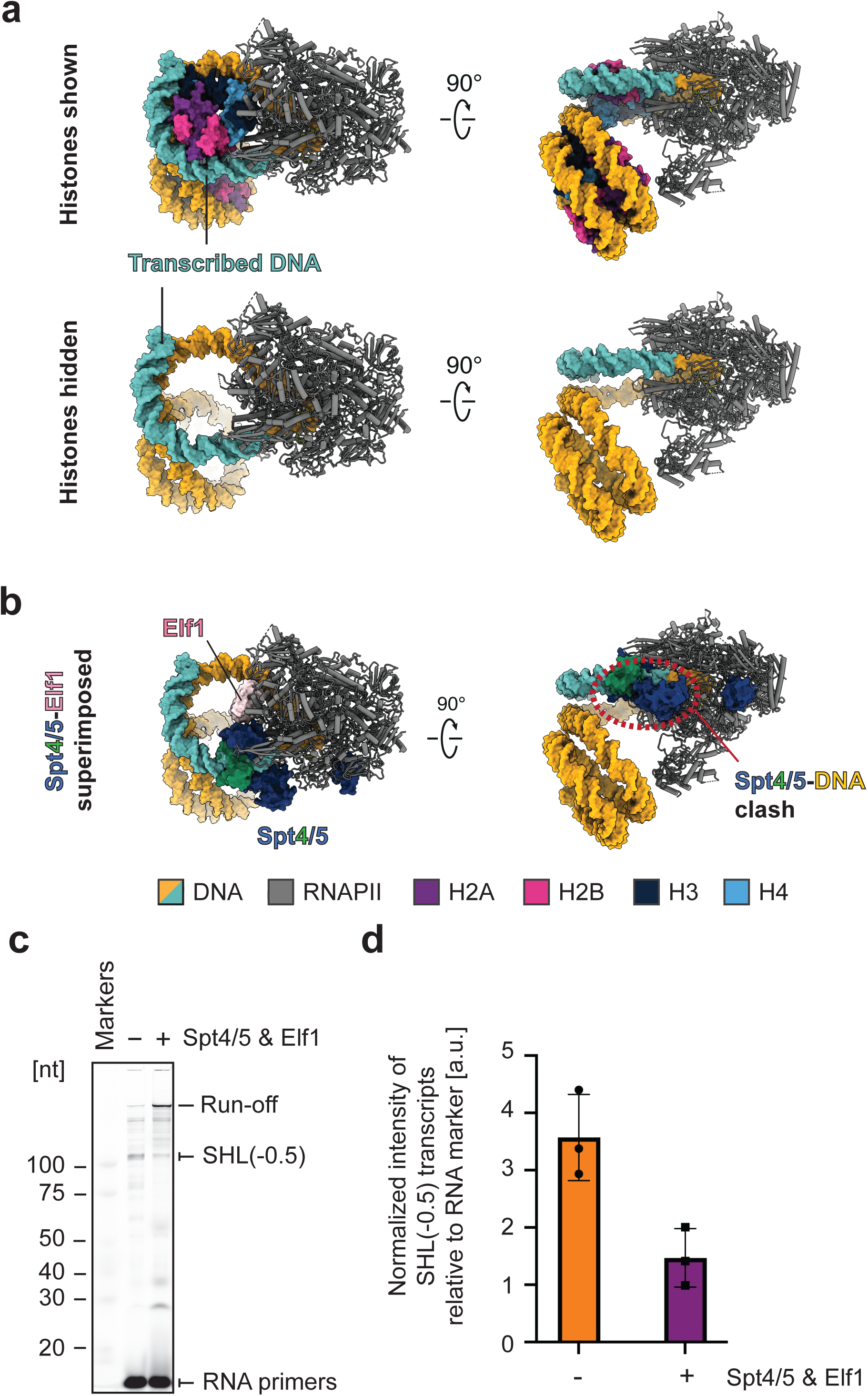
Upstream DNA loops back onto the hexasome subunit in the RNAPII-OLDN^SHL(−0.5)^ complex. **a,** *Upper panel:* Transcribed upstream DNA (cyan) loops back and rewraps onto the hexasome subunit of the OLDN. *Lower panel:* Histones are hidden to allow clearer visualization of the DNA trajectory. **b,** Superposition of transcription factors Spt4/5 (blue-green) and Elf1 (light pink) (PDB: 6IR9) onto the RNAPII–OLDN^SHL(−0.5)^ structure reveals steric clashes with the upstream DNA loop. **c,** Representative transcription assays of the hexasome–octasome OLDN configuration in the absence and presence of transcription factors Spt4/5 and Elf1. Transcription experiments were performed in three independent replicates. **d,** Quantification of the SHL(−0.5) transcripts from **c**. RNA transcript intensities were normalized to a reference ladder band on the same gel. Error bars indicate standard deviation (n=3).

In this complex, the upstream DNA region is rewrapped in the H2A-H2B surface exposed by the DNA peeling mediated by the RNAPII progression (Fig. 3a). The upstream DNA looping has also been observed in the canonical nucleosome transcription by the RNAPII paused at the SHL(-1) position^21^. However, the DNA looping in the OLDN has a substantially different structure from that in canonical nucleosome. In the RNAPII-OLDN^SHL(−0.5)^ structure, hexasome moiety directly contacts with the clamp head of the RNAPII, and is deeply incorporated into the RNAPII cleft (Fig. 3a). Consequently, the length of DNA between RNAPII and the histones in the hexasome disk is reduced compared with that in the canonical nucleosome. In the canonical nucleosome, the clamp head of RNAPII directly contacts with nucleosomal DNA at ∼SHL(+5.5) position, which is wrapped around the distal H2A-H2B dimer.

Since this distal H2A-H2B dimer is absent in OLDN, RNAPII associateed more tightly with histones and formed a different DNA looping structure with the transcribed upstream DNA region in OLDN.

In the previous study, we found that the template DNA looping of the RNAPII-nucleosome complex occurs in the absence of transcription elongation factors, Spt4/5 and Elf1^21^. Since Spt4/5 and Elf1 are reported to bind near the DNA exit site of RNAPII and inhibit the DNA bending required for the DNA looping formation, the DNA looping may not occur in the presence of Spt4/5 and Elf1^21,22^ (Fig. 3b). Consistently, the SHL(−0.5) pausing was drastically reduced when elongation factors, Spt4/5 and Elf1, were added to the reaction mixture of the OLDN transcription by RNAPII (Fig.3c, d).

## Discussion

Our study uncovers a previously unrecognized directional asymmetry in how RNAPII engages and traverses the overlapping dinucleosome (OLDN). Although OLDNs accumulate immediately downstream of transcription start sites^11,13,14^, how RNAPII negotiates this compact and asymmetric structure remained unclear. By combining *in vitro* transcription with cryo-EM, we show that the hexasome side of the OLDN constitutes a permissive entry point for RNAPII, whereas the octasome side imposes a substantially higher barrier (Fig.4). This directional effect positions the OLDN not merely as a passive chromatin obstacle but as an intrinsically polarity-sensitive regulator of early transcription elongation.

**Figure 4.**
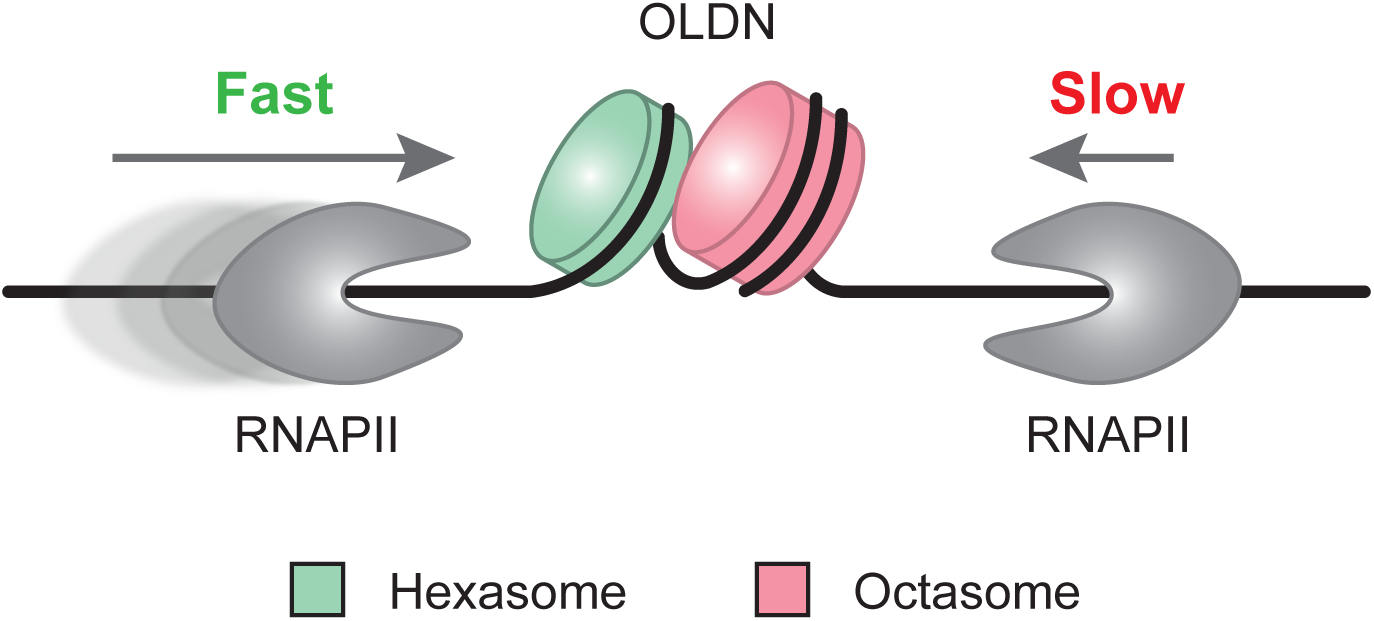
Model for direction-dependent transcriptional regulation by OLDN configuration. OLDNs may adopt distinct configurations in the genome. When RNAPII approaches the OLDN from the hexasome side, transcription proceeds more productively. In contrast, approach from the octasome side results in markedly reduced transcriptional output. This direction-dependent behavior suggests a potential mechanism by which OLDN configuration modulates RNAPII transcription.

A central finding is the emergence of a distinctive pausing event when RNAPII approaches the hexasomal SHL(−0.5) position. The cryo-EM structure at this native pause reveals extensive remodeling of the dinucleosome: the hexasome and octasome adopt an almost perpendicular configuration, with RNAPII simultaneously engaging both subnucleosomal units. This remodeling is driven by direct contacts of the RNAPII clamp with the hexasome and of the Rpb4/7 stalk with the octasome (Fig.2), effectively opening the interface between the two nucleosomes. These observations indicate that RNAPII is not a passive traveler but an active mechanical remodeler of the OLDN, reshaping its architecture to facilitate progression.

The RNAPII-OLDN^SHL(−0.5)^ structure further reveals a prominent upstream DNA loop re-engaging the remaining H2A-H2B dimer of the hexasome. The upstream ∼40-bp DNA loop is similar to the DNA loop formed at SHL(-1) in a canonical nucleosome^21^. In the OLDN, however, the absence of one H2A-H2B dimer between the hexasome and the octasome unit brings the hexasome disc closer to RNAPII. This facilitates the upstream transcribed DNA to capture the exposed histone surface on the hexasome more effectively, thereby promoting efficient loop formation. Importantly, elongation factors Spt4/5 and Elf1, which restrict DNA bending at the RNAPII exit site, strongly suppress the SHL(−0.5) pausing, indicating that loop formation promoted in OLDN is a regulated, factor-sensitive intermediate during traversal.

Following traversal of the hexasome, RNAPII must engage the adjacent octasome. Although no linker DNA separates the two moieties, the presence of a hexasome may alter the local histone-DNA constraints at the octasome entry region, thereby modulating the effective transcriptional barrier, contributing to a higher overall transcription efficiency for OLDN with a hexasome-octasome configuration.

The directional behavior of RNAPII on OLDNs has important implications for transcription start site proximal regulation. Because OLDNs decorate the +1 region, their structural asymmetry may contribute to enforcing transcriptional directionality: RNAPII entering in the productive (sense) direction encounters a hexasome first, whereas antisense transcription would face a more rigid octasome barrier. Such polarity could help suppress bidirectional transcription and sharpen promoter output. Moreover, because OLDNs can be generated by chromatin remodelers and may also arise as intermediates during histone eviction and redeposition, RNAPII-mediated remodeling of OLDNs is likely to be tightly integrated with multiple chromatin-dependent pathways that regulate promoter-proximal transcription.

Our reconstituted system isolates the mechanical features of RNAPII-OLDN interactions, but additional layers of regulation likely operate *in vivo*. Remodelers, histone chaperones, histone variants, and histone modifications may shift the stability of OLDN and/or modulate the magnitude of the directional asymmetry. Together, these results establish the OLDN as a dynamic, directionally tuned chromatin architecture that actively shapes early transcription elongation. The structural insights presented here provide a framework for understanding how non-canonical nucleosome states influence transcriptional regulation *in vivo*.

## Methods

### Preparation of proteins

Human histones H2A, H2B, H3.3, H4 and histone octamers were prepared as described previously^23^. In brief, the coding sequences of H2A, H2B, H3.3, and H4 were inserted into pET15b vectors (Novagen) individually. The His_6_-tagged histones were expressed in the *E. coli* BL21 (DE3) strain (H2A, H2B and H3.3) or the JM109 (DE3) strain (H4), and purified by Ni-NTA (Qiagen) affinity column chromatography under denaturing conditions. The His_6_-tags were later cleaved by thrombin protease (Wako), and the histones were purified by MonoS cation-exchange column chromatography (Cytiva).

The purified histones were dialyzed against water and lyophilized.

Histone octamers were generated by mixing H2A, H2B, H3.3, and H4 with an equal molar amount, followed by dialysis against refolding buffer (10mM Tris-HCl (pH 7.5), 1mM EDTA, 2M NaCl, and 5mM 2-mercaptoethanol). The sample was further purified through a Superdex 200 gel filtration column (Cytiva). The purified histone octamers were then flash-frozen and stored at -80°C until use.

RNAPII was purified from a genetically modified strain of *Komagataella phaffii*, containing a TAP tag in the Rpb2 subunit, as described before^18^. Elongation factors were prepared using *E. coli* expression system also as previously described^18^. Briefly, DNA fragments encoding *K. phaffii* TFIIS, Spt4/5, and Elf1 were cloned into the pET-47b vector (Novagen) and transformed *E. coli* strain KRX (Promega). Proteins were expressed with an N-terminal His-tag, which was subsequently removed by HRV-3C protease. Purified *K. phaffii* RNAPII, TFIIS, Spt4/5 and Elf1 were all dialyzed against KpRNAPII buffer (20 mM HEPES-KOH (pH 7.5), 150 mM potassium acetate, 1 μM zinc acetate, 0.1 mM tris(2-carboxyethyl)phosphine (TCEP) and 5% glycerol) and stored at -80°C until use.

### Preparation of the OLDN for the transcription assay (biochemical analysis)

For biochemical analysis, OLDN (hexasome-octasome or octasome-hexasome configurations) with a total length of 323 bp were prepared. Similar to previously reported methods for preparing the nucleosome template for transcription^19^, the OLDN templates for transcription are prepared in a two-step manner. The “OLDN core” is first reconstituted to ensure appropriate histone positioning. This is then followed by the ligation of a DNA fragment that contains the mismatched bubble DNA for transcription initiation, on either side of the “OLDN core” to form the hexasome-octasome or octasome-hexasome configuration.

In our previous study, we designed a 250 bp Widom 601-based sequence^24^ that could facilitate the formation of the OLDN in a fixed direction, with the hexasome subunit at one side and the octasome subunit at the other^13^. In this study, we utilized this knowledge and extended the OLDN DNA sequence on either hexasome or octasome side to include a BglI digested sticky end for subsequent ligation reactions. Specifically, the DNA fragments containing 250 bp Widom sequence and a short segment harbouring a BglI recognition site (either on the hexasome side or octasome side) were amplified by PCR and digested by BglI (Takara). The digested DNA fragments were purified by phenol chloroform extraction and ethanol precipitation, and further purified by the Prep Cell apparatus (Bio-Rad). The resulting DNA fragment containing the Widom 601 DNA sequence is as follows:

For OLDN (hexasome-octasome configuration), the non-template strand is as (258 bp): 5’- TGGCC GTTTT CGTGT TTCCC GGTGC CGTGG CCGCT CTTTT GGTCG TTGTC AGCTC TAGCA CCGCT TAAAC GCACG TACGC GCTGT CCCCC GCGTT TTAAC CGCCA AGGGG ATTAC TCCCT AGTCT CCAGG CTCGA GCTCA ATTGG TCGTA GACAG CTCTA GCACC GCTTA AACGC ACGTA CGCGC TGTCC CCCGC GTTTT AACCG CCAAG GGGAT TACTC CCTAG TCTCC AGGCA CGTGT CAGAT ATATA CATCC GAT - 3’; the template strand is as (261 bp): 5’-ATCGG ATGTA TATAT CTGAC ACGTG CCTGG AGACT AGGGA GTAAT CCCCT TGGCG GTTAA AACGC GGGGG ACAGC GCGTA CGTGC GTTTA AGCGG TGCTA GAGCT GTCTA CGACC AATTG AGCTC GAGCC TGGAG ACTAG GGAGT AATCC CCTTG GCGGT TAAAA CGCGG GGGAC AGCGC GTACG TGCGT TTAAG CGGTG CTAGA GCTGA CAACG ACCAA AAGAG CGGCC ACGGC ACCGG GAAAC ACGAA AACGG CCACC A -3’

For OLDN (octasome-hexasome configuration): the non-template strand is as (258 bp): 5’-TGGCG CAAAT CGGAT GTATA TATCT GACAC GTGCC TGGAG ACTAG GGAGT AATCC CCTTG GCGGT TAAAA CGCGG GGGAC AGCGC GTACG TGCGT TTAAG CGGTG CTAGA GCTGT CTACG ACCAA TTGAG CTCGA GCCTG GAGAC TAGGG AGTAA TCCCC TTGGC GGTTA AAACG CGGGG GACAG CGCGT ACGTG CGTTT AAGCG GTGCT AGAGC TGACA ACGAC CAAAA GAGCG GCCAC GGCAC CGGGA AACAC GAT -3’; the template strand is as (261 bp): 5’ – ATCGT GTTTC CCGGT GCCGT GGCCG CTCTT TTGGT CGTTG TCAGC TCTAG CACCG CTTAA ACGCA CGTAC GCGCT GTCCC CCGCG TTTTA ACCGC CAAGG GGATT ACTCC CTAGT CTCCA GGCTC GAGCT CAATT GGTCG TAGAC AGCTC TAGCA CCGCT TAAAC GCACG TACGC GCTGT CCCCC GCGTT TTAAC CGCCA AGGGG ATTAC TCCCT AGTCT CCAGG CACGT GTCAG ATATA TACAT CCGAT TTGCG CCACC A-3’.

Each fragment was then mixed with the histone octamer and subjected to salt dialysis to form the “OLDN core”. After salt dialysis, the OLDN cores were then purified by the Prep Cell apparatus (Bio-Rad). A DY647-labeled DNA fragment that contains the bubble DNA was ligated onto the BglI sticky end of the OLDN template by T4 ligase (NIPPON GENE).

The DNA fragment that contains the mismatched bubble is as follows: Non-template strand (65 bp) 5’-GCTTA CGTCA GTC TGGCCATCT TTGTG TTTGG TGTGT TTGGG GACAA ACACG TCCGG AATGA TGG -3’; Template strand (62bp) 5’-TCATT CCGGA CGTGT TTGTC CCCAA ACACA CCAAA CACAA GAGCTAATT GACTG ACGTA AGC -3’. The RNA primer for transcription initiation is as: 5’-AUAAUUAGCUC-3’.

The final OLDN DNA constructs are summarized in Table 1.

**Table 1.**
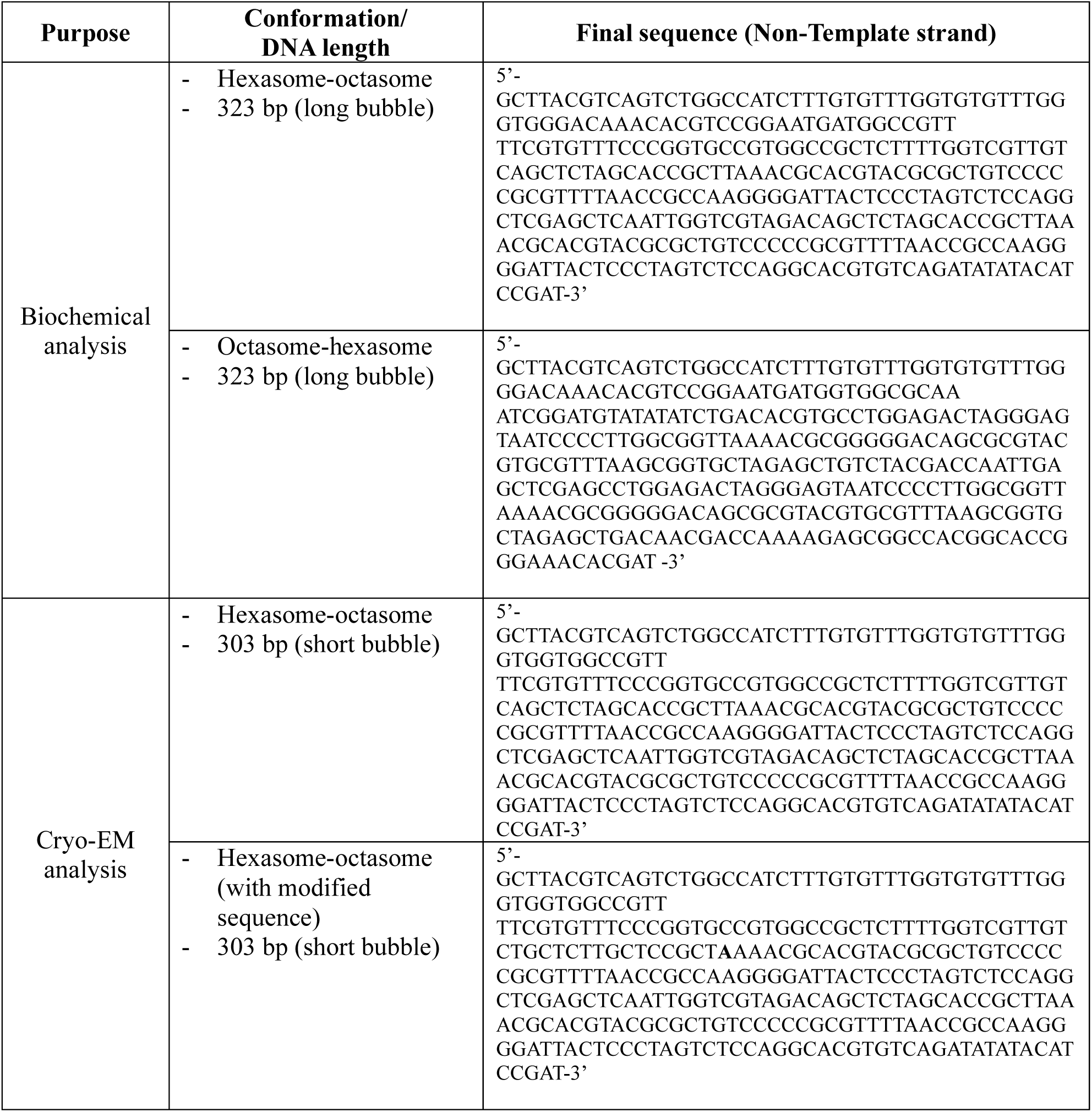
The complete OLDN DNA sequences used in this study.

### *In vitro* transcription assays

The transcription assays conducted in a time course. 0.1 μM OLDN, 0.3 μM *K. phaffii* RNAPII, 0.6 μM *K. phaffii* TFIIS, and 0.2 μM DY647-labeled primer RNA (5′-DY647-AUAAUUAGCUC-3′, Dharmacon) were mixed in 10 µL of mixture that contained 30 mM HEPES-KOH (pH 7.5), 5 mM MgCl_2_, 80 mM potassium acetate, 0.4 μM zinc acetate, 40 μM Tris(2-carboxyethyl) phosphine, 0.1 mM DTT, 2.5% glycerol, 0.4 mM UTP, CTP, GTP, and ATP. The reaction mixture was incubated at 30 °C and 2 μL of reaction aliquot was collected after 2, 5, 10 and 30 mins. The reaction was quenched by adding 1 μL of Proteinase K mixture (0.5 mg/mL Proteinase K (Roche), 200 mM Tris-HCl, 80mM EDTA) and further incubated for 10 mins at room temperature.

Subsequently, 12 μL of Hi-Di formamide (Thermo Fisher Scientific) was added to the aliquot and thermally denatured at 95 °C for 5 mins.

The transcription assays conducted to investigate the effect of Spt4/5 and Elf (Fig. 3c) have a modified reaction condition. 0.1 μM OLDN, 0.3 μM *K. phaffii* RNAPII, 0.6 μM *K. phaffii* TFIIS, 1.2 μM *K. phaffii* Spt4/5, 3 μM *K. phaffii* Elf1 and 0.2 μM DY647-labeled primer RNA (5′-DY647- AUAAUUAGCUC-3′, Dharmacon) were mixed in 10 µL of mixture that contained 30 mM HEPES-KOH (pH 7.5), 5 mM MgCl_2_, 80 mM potassium acetate, 0.4 μM zinc acetate, 40 μM Tris(2-carboxyethyl) phosphine, 0.1 mM DTT, 2.5% glycerol, 0.4 mM UTP, CTP, GTP, and ATP. The reaction mixture was incubated at 30 °C for 10 mins and 2 μL aliquot was taken. The aliquot reaction was quenched by adding 1 μL of Proteinase K mixture (0.5 mg/mL Proteinase K (Roche), 200 mM Tris-HCl, 80mM EDTA) and further incubated for 10 mins at room temperature. Subsequently, 12 μL of Hi-Di formamide (Thermo Fisher Scientific) was added to the aliquot and thermally denatured at 95 °C for 5 mins.

In all transcription assays, the length of elongated RNA products was assessed by 10% urea denaturing polyacrylamide gel electrophoresis (Urea denaturing-PAGE). The DY467-labeled RNA primer enables the detection of RNA products using the Amersham Typhoon scanner (Cytiva) in the Cy5 channel. The band intensities of the transcripts were quantified and normalized with reference ladder bands on the same gel, using the ImageQuant™ TL software (GE Healthcare).

### Preparation of the OLDN for the structural analyses

For cryo-EM analysis, OLDNs with the hexasome-octasome configuration were used. Basically, the OLDN preparation method was carried over from previous section.

However, to improve sample homogeneity, a shorter linker DNA was used (Non-template strand (45 bp) 5’-GCTTA CGTCA GTCTG GCCAT CTTTG TGTTT GGTGT GTTTGG GTGG -3’; Template strand (42 bp) 5’- CCCAA ACACA CCAAA CACAA GAGCT AATTG ACTGA CGTAA GC -3’). This results in a 303 bp construct, which is 20 bp shorter than the construct used for biochemical analysis. The successful reconstitution of this OLDN is confirmed by both native PAGE and SDS-PAGE, and the OLDN construct was used for solving the RNAPII-OLDN^SHL(−6)^ structure .

To solve the RNAPII-OLDN^SHL(−0.5)^ structure, the OLDN core sequence was modified, so that when combined with 3’-dATP, UTP, CTP, and GTP, RNAPII would pause around hexasome SHL(−0.5). The modified OLDN core sequence is as follows (the altered bases were underlined, and the engineered pausing site is in **bold**): the non-template strand is as (258 bp.): 5’- TGGCC GTTTT CGTGT TTCCC GGTGC CGTGG CCGCT CTTTT GGTCG TTGTC <u>T</u>GCTC T<u>T</u>GC<u>T</u> CCGCT **<u>A</u>**AAAC GCACG TACGC GCTGT CCCCC GCGTT TTAAC CGCCA AGGGG ATTAC TCCCT AGTCT CCAGG CTCGA GCTCA ATTGG TCGTA GACAG CTCTA GCACC GCTTA AACGC ACGTA CGCGC TGTCC CCCGC GTTTT AACCG CCAAG GGGAT TACTC CCTAG TCTCC AGGCA CGTGT CAGAT ATATA CATCC GAT - 3’; the template strand is as (261 bp): 5’-ATCGG ATGTA TATAT CTGAC ACGTG CCTGG AGACT AGGGA GTAAT CCCCT TGGCG GTTAA AACGC GGGGG ACAGC GCGTA CGTGC GTTTA AGCGG TGCTA GAGCT GTCTA CGACC AATTG AGCTC GAGCC TGGAG ACTAG GGAGT AATCC CCTTG GCGGT TAAAA CGCGG GGGAC AGCGC GTACG TGCGT TT**<u>T</u>**AG CGG<u>A</u>G C<u>A</u>AGA GC<u>A</u>GA CAACG ACCAA AAGAG CGGCC ACGGC ACCGG GAAAC ACGAA AACGG CCACC A -3’. The successful reconstitution of this OLDN is confirmed by both native PAGE and SDS-PAGE, and the OLDN construct was used for solving the RNAPII-OLDN^SHL(−6)^ structure.

### Cryo-EM sample preparation

The workflow for RNAPII-OLDN complexes preparation is composed of three major steps: preparation of the transcription reaction mixture, Gradient-Fixation (GraFix), and grid preparation. As mentioned in the previous section, we prepared two samples and resolved the RNAPII-OLDN^SHL(−6)^ complex and the RNAPII-OLDN^SHL(−0.5)^ complex from each sample, respectively.

For the preparation of the RNAPII-OLDN^SHL(−6)^ complex, 0.1 μM 303 bp OLDN, 0.3 μM *K. phaffii* RNAPII, 0.3 μM *K. phaffii* TFIIS, and 0.2 μM DY647-labeled primer RNA (5′-Dy647- AUAAUUAGCUC-3′, Dharmacon) was mixed in 650 µL of reaction mixture (26 mM HEPES-KOH (pH 7.5), 5 mM MgCl_2_, 50 mM potassium acetate, 0.2 μM zinc acetate, 20 μM Tris(2-carboxyethyl) phosphine, 0.1 mM DTT, 1.5% glycerol, 0.4 mM UTP, CTP, GTP, and ATP). The reaction mixture was incubated at 30°C for 20 mins, and was subsequently quenched by the addition of EDTA (final concentration: 50 mM).

For the preparation of the RNAPII-OLDN ^SHL(−0.5)^ complex, 0.1 μM 303 bp OLDN with defined pausing site, 0.1 μM *K. phaffii* RNAPII, 0.1 μM *K. phaffii* TFIIS, and 0.4 μM DY647-labeled primer RNA (5′-Dy647- AUAAUUAGCUC-3′, Dharmacon) was mixed in 1080 µL of reaction mixture (26 mM HEPES-KOH (pH 7.5), 5 mM MgCl_2_, 50 mM potassium acetate, 0.2 μM zinc acetate, 20 μM Tris(2-carboxyethyl) phosphine, 0.1 mM DTT, 1.5% glycerol, 0.4 mM UTP, CTP, GTP, and 3’-dATP). The use of 3’-dATP can pause RNAPII at the defined position. The reaction mixture was incubated at 30°C for 30 mins then quenched by the addition of EDTA (final concentration: 50 mM).

To further stabilize and purify the RNAPII-OLDN complexes, the reaction mixture were subjected to GraFix^25^ as previously described^19^. The collected fractions were dialyzed against a buffer that contains 20mM HEPES-KOH (pH 7.5), 50 mM potassium acetate, 0.2 μM zinc acetate, 0.1mM Tris(2-carboxyethyl) phosphine.

Subsequently, the RNAPII-OLDN^SHL(−6)^ and RNAPII-OLDN^SHL(−0.5)^ complexes were concentrated to 136.1 ng/µL and 143.7 ng/µL (DNA concentration), respectively, using an Amicon Ultra 100K centrifugal filter unit (Merck). Finally, the RNA composition of sample was confirmed by urea-denaturing PAGE.

Immediately before application onto the cryo-EM grid, fresh detergent Tween-20 was added to the sample solution to a final concentration of 0.0025% for optimizing the particle quality and integrity^26^. R1.2/1.3, Cu, 200 mesh Quantifoil grids (Qunatifoil Micro Tools) were glow discharged by PIB-10 Ion Bombarder (Vacuum Device Inc. Japan). 2 µL of samples were loaded onto cryo-EM grids, followed by plunge-freezing in liquid ethane with a Vitrobot Mark IV (4°C and 100% humidity) (Thermo Fisher Scientific).

### Data acquisition

Cryo-EM images were collected by a Krios G4 transmission electron microscope (Thermo Fisher Scientific), equipped with a K3 direct electron detector (Gatan). The automated data acquisition was executed by EPU software (Thermo Fisher Scientific) and operated at 300kV at a pixel size of 1.06Å. Each image was fractionated into 40 frames and saved as non-gain-normalized compressed TIFF files.

### Cryo-EM image processing of the RNAPII-OLDN^SHL(−6)^ complex

Cryo-EM movies from the RNAPII-OLDN^SHL(−6)^ datasets were imported into CryoSPARC^27^ v4 for single-particle analysis. Patch Motion Correction and Patch CTF Estimation were performed, after which low-quality micrographs were manually removed using the Curate Exposures tool.

A trained Topaz model, generated based on a subset of 1000 selected micrographs from the full datasets, was subsequently applied to the full datasets. Picked particles were extracted and subjected to two rounds of 2D classification. After the first round, classes corresponding to RNAPII dimers lacking OLDN density were identified and removed from further analysis. The second round of 2D classification yielded a cleaner particle set enriched for RNAPII-OLDN complexes.

Selected particles were then subjected to two rounds of heterogeneous refinement. An additional round of 2D classification was performed to remove a small number of remaining junk particles. Local motion correction was then applied, followed by another round of heterogeneous refinement to select the class exhibiting the clearest OLDN^SHL(−6)^ density. After recentering, particles were re-extracted and refined using homogeneous refinement. Finally, local motion correction, homogeneous refinement and 3D flexible refinement were performed to yield the final map.

### Cryo-EM image processing of the RNAPII-OLDN^SHL(−0.5)^ complex

Cryo-EM movies from the RNAPII-OLDN^SHL(−0.5)^ dataset were imported into CryoSPARC^27^ v4 for single-particle analysis. Patch Motion Correction and Patch CTF Estimation were performed, after which low-quality micrographs were manually removed using the Curate Exposures tool.

A trained Topaz model, generated from an initial analysis of the dataset, was applied to pick particles from the full dataset. Picked particles were extracted and subjected to two rounds of 2D classification. Subsequently, three rounds of heterogeneous refinement were performed, and particles exhibiting clear RNAPII and OLDN density were selected. The selected particles were recentered, re-extracted, and refined using homogeneous refinement.

To improve the density of the OLDN, 3D classification was performed using a focus mask on the octasome subunit within the OLDN. Selected classes were then subjected to local motion correction followed by homogeneous refinement. 3D flexible refinement was performed to yield the final overall map.

To aid model building, local maps focused on either the octasome or the RNAPII were also generated.

### Model building

The RNAPII–OLDN^SHL(−6)^ model was generated using the atomic model of the *K. phaffii* RNAPII elongation complex stalled at SHL(–6) on a canonical nucleosome^19^ (PDB: 6A5O) and the overlapping trinucleosome (OLTN)^14^ (PDB: 8IHL). The assembled structure was refined against the RNAPII–OLDN^SHL(−6)^ density map using the phenix.real_space_refine tool in the Phenix package^28^, followed by manual inspection and adjustment in the Coot software^29,30^.

The RNAPII–OLDN^SHL(−0.5)^ model was generated using the atomic model of the *K. phaffii* RNAPII elongation complex^9^ (PDB: 7XSE) and the OLTN^14^ (PDB: 8IHL). Focused refinement maps of the octasome-DNA and RNAPII-hexasome regions were used to refine the corresponding components. The complete model was then refined against the RNAPII–OLDN^SHL(−0.5)^ overall density map using phenix.real_space_refine tool in the Phenix package^28^, followed by manual inspection and adjustment in the Coot software^29,30^.

In both structures, the histone sequences of the input models were modified to match the histones that were used in this study. All structural figures were prepared using UCSF ChimeraX^31^.

### Use of generative AI

During the preparation of this work, the authors used ChatGPT, a Large Language Model developed by OpenAI, in a strictly auxiliary role to support the writing process. This included assistance with phrasing and grammar correction. After using ChatGPT, the authors reviewed and edited the content as needed and take full responsibility for the content of the published article.

### Data availability

The data that support this study are available from the corresponding authors upon reasonable request. The cryo-EM reconstructions and atomic models of the RNAPII-OLDN^SHL(−0.5)^ complex and the RNAPII-OLDN^SHL(−6)^ complex generated in this study have been deposited in the Electron Microscopy Data Bank and the Protein Data Bank, under the accession codes EMD-XXXXX and PDB ID XXXX, and EMD-XXXXX and PDB ID XXXX, respectively. The structures used in this study can be found in the Protein Data Bank under the accession codes: 6A5O, 7XSE and 8IHL.

## Acknowledgements

We would like to express our deepest gratitude to M. Dacher, and Y. Takeda (Univ. of Tokyo) for their assistance. This work was supported in part by JSPS KAKENHI grant number JP25K18403 [to C.H.H.], JSPS KAKENHI Grant Numbers JP20H05690 [to S.S., T.K.], JP24H00062 [to S.S., T.K.], JP20H03201 [to H.E., T.K.], JP23K17392 [to T.K.], JP24H02319 [to H.K.], JP24H02328 [to H.K.], JP23H05475 [to H.K.], JST ERATO Grant Number JPMJER1901 [to H.K.], CREST JPMJCR24T3 [to H.K.], AMED Grant Number JP25ama121002 [to Y.T.], and AMED BINDS Grant Number JP25ama121009 [to H.K.].

## Author contributions

Z.C., and H.T., prepared the RNAPII-OLDN complexes, performed biochemical analysis. Z.C., H.T., and C.-H.H. collected cryo-EM data. C.-H.H., Z.C., M.O, and Y.T. performed cryo-EM analysis. H.E. and S.S. prepared RNAPII, TFIIS, Spt4/5 and Elf1. H.K. conceived, designed, and supervised all of the work. Z.C., C.-H.H. and H.K. prepared all figures and wrote the paper. All of the authors discussed the results and commented on the manuscript.

